# Deep learning-based image quantification of epithelial cell shapes and its application to polycystic kidney disease

**DOI:** 10.1101/2025.08.13.670046

**Authors:** Johannes Jahn, Alexis Hofherr, Clara Consoli, Berenike Fajen, Rebekka Goll, Greta Theresa Liedtke, Adrian Boehm, Friederike Selbach, Paul Christoph Zeisler, Vanessa Weichselberger, Anne-Kathrin Classen, Lukas Westermann, Tilman Busch, Michael Köttgen

## Abstract

Cell shape is a fundamental determinant of tissue architecture and organ function. In epithelial tissues, cytoskeletal organization and tight junctions regulate cell geometry, shaping functional tissue units. Disruption of these mechanisms may cause diseases such as autosomal dominant polycystic kidney disease (ADPKD), in which cyst formation is characterized by abnormal regulation of epithelial cell shape. The mechanisms of cystogenesis remain incompletely understood, highlighting the need for robust, high-throughput methods to quantify the morphology of epithelial cells. Here, we present a fully automated, deep learning-based image analysis pipeline to quantify epithelial cell shape and tight junction morphology from immunofluorescence images. Our approach employs a U-Net convolutional neural network for accurate segmentation of fluorescence labeled tight junctions. We introduce novel algorithms to quantify overall cell shape and tight junction morphology, as well as to estimate cytoskeletal traction at shared cell borders. Our analysis pipeline objectively identifies subtle morphogenetic changes associated with disease-related mutations, applied to a genetically modified Madin-Darby Canine Kidney cell model of ADPKD. The method enables high-throughput, standardized analysis, reduces observer bias, and facilitates comparison across experiments. We further demonstrate the pipeline’s generalizability by applying it to Drosophila egg chamber epithelia. Our results establish a robust and scalable framework for analyzing cell shape and mechanical interactions in epithelial tissues, with broad applications in phenotypic screening, disease modeling, and morphogenesis research.

**Author Summary:** The shape of epithelial cells is critical for organ function. In the kidney, properly shaped epithelial cells assemble to tubules ensuring efficient waste excretion as well as body electrolyte and water balance. Disruption of cell shape regulation can lead to diseases such as autosomal dominant polycystic kidney disease (ADPKD), characterized by cyst formation and displacement of normal kidney tissue. Traditionally, analysis of epithelial cell morphology has relied on manual, low-throughput methods, which are time-consuming and prone to error. To overcome these limitations, we developed a fully automated, artificial intelligence-based pipeline that rapidly and reliably quantifies cell shape and junctional organization from microscopic images. We validated our approach using a cellular model of ADPKD, demonstrating clear differences in cell shape and junctional structure between normal and mutant cells harboring mutations in PKD-related genes. Our method enables efficient, objective analysis of large datasets and provides a powerful tool for understanding the mechanisms underlying cell shape regulation in health and disease.

## Introduction

Epithelial cells are the fundamental building blocks of many organs, and their ability to adopt specific shapes underlies critical morphogenetic processes during development [1]. Cell shape changes, including apical constriction, drive tissue folding, tubulogenesis, and lumen formation [2]. These processes are powered by the actomyosin cytoskeleton and coordinated through cell–cell junctions, particularly adherens and tight junctions [3].

At the organ level, the geometry of epithelial tissues directly contributes to organ shape and function. In the kidney, precise control of epithelial tubule geometry is crucial for proper organ function [4]. Disruption of the cellular mechanisms controlling epithelial morphology can lead to severe developmental disorders and diseases, including Autosomal Dominant Polycystic Kidney Disease (ADPKD), a monogenic disorder caused by mutations in *PKD1* and *PKD2* [5,6]. These genes encode proteins involved in signal transduction in primary cilia, and their dysfunction leads to altered cytoskeletal regulation, dedifferentiation, and cyst formation [7–13].

Quantitative analysis of epithelial morphology *in vitro* can provide valuable mechanistic insights. However, current methods often rely on manual annotation or low-throughput approaches, limiting scalability and reproducibility. Recent advances in deep learning, particularly convolutional neural networks such as U-Net, have transformed biomedical image classification and segmentation, enabling exceptional performance even with limited datasets [14–16]. This facilitates automated and standardized quantification of cell shape across large datasets [17].

Here, we present a deep learning-based workflow for automated analysis of epithelial morphology, with a focus on tight junction staining using Zonula occludens 1 (ZO1) as a marker. We apply this pipeline to a genetically modified Madin-Darby Canine Kidney (MDCK) cell model of ADPKD and introduce novel quantitative features to describe tight junction configuration and cytoskeletal traction. Our results demonstrate how this system enables objective, high-throughput phenotyping and reveals subtle morphogenetic differences associated with disease.

## Results

### Deep learning-based segmentation of ZO1 images

Forces generated by the cytoskeleton are essential for cell and organ shape during development, homeostasis and regeneration [18,19]. To investigate cell shapes in polarized kidney epithelial cells, we imaged the tight-junction-associated protein ZO1. Immunofluorescence staining of ZO1 revealed distinct tight junction patterns in wild-type (WT) and *PKD2*^*-/-*^ MDCK cells. WT cells showed heterogeneous, meandering tight junctions with a pronounced riffled pattern, whereas *PKD2*^*-/-*^ cells displayed smoother, more uniform junctions (Fig 1). Re-expression of PKD2 partially rescued this phenotype by restoring the characteristic riffled pattern (D in S1 Fig). PKD1-/- cells also displayed this specific phenotype (B in S1 Fig.).

**Fig 1.**
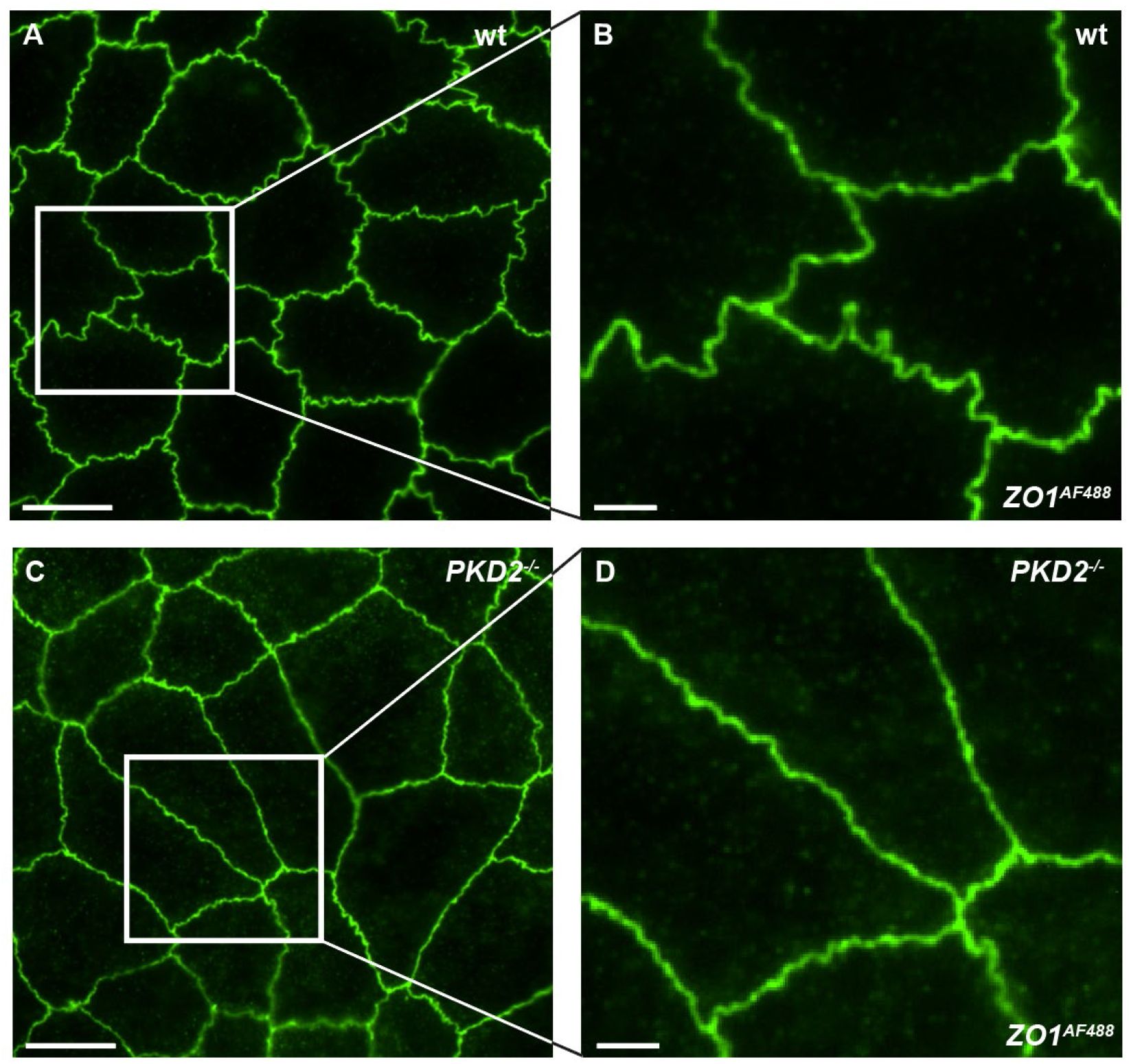
Tight junction meandering as a morphological parameter. Representative immunofluorescence images of MDCK WT and *PKD2*^*–/–*^ cells. Cells were stained with an α-Zonula occludens 1 primary antibody and an Alexa Fluor 488-conjugated secondary antibody. (A) WT cells show prominent membrane riffles, characterized by indentations and a meandering junctional pattern. (B) Enlarged view of panel (A). (C) *PKD2*^*–/–*^ cells exhibit smooth cell borders with minimal ruffling lacking major indentations. (D) Enlarged view of panel (C). Scale bars: 10 μm (A, C); 2.5 μm (B, D). MDCK = Madin-Darby Canine Kidney; *PKD2* = Polycystic kidney disease gene 2; WT = Wildtype, ZO1 = Zonula occludens protein 1.

Based on these findings, we developed an end-to-end processing workflow that translates acquired images into quantifiable values, including tight junction morphology (Fig 2). To address the limitations of conventional threshold-based segmentation methods, we trained a deep learning segmentation system using a U-Net convolutional network to detect ZO1-stained tight junctions (S2 Fig). Training images were randomly selected from diverse experiments to minimize selection bias and to improve generalizability. The training dataset included not only images with varying signal intensities but also with considerable background noise to enhance model robustness. Labeling of the tight junctions followed a predefined strategy, excluding out-of-focus regions to avoid interference with downstream analysis. Segmentation quality, evaluated by multiple researchers, showed consistently accurate detection across varying background signals and artifacts, achieving an intersection over union of 0.771 after 10.000 iterations. Deviations from the ground truth were small, typically within one pixel, and had only negligible effect on overall morphology and the characteristic meandering junctional pattern. Our setup processed approximately 1,000 images per hour (1388×1040 px; ∼0.015 mm^2^), enabling efficient high-throughput analysis.

**Fig 2.**
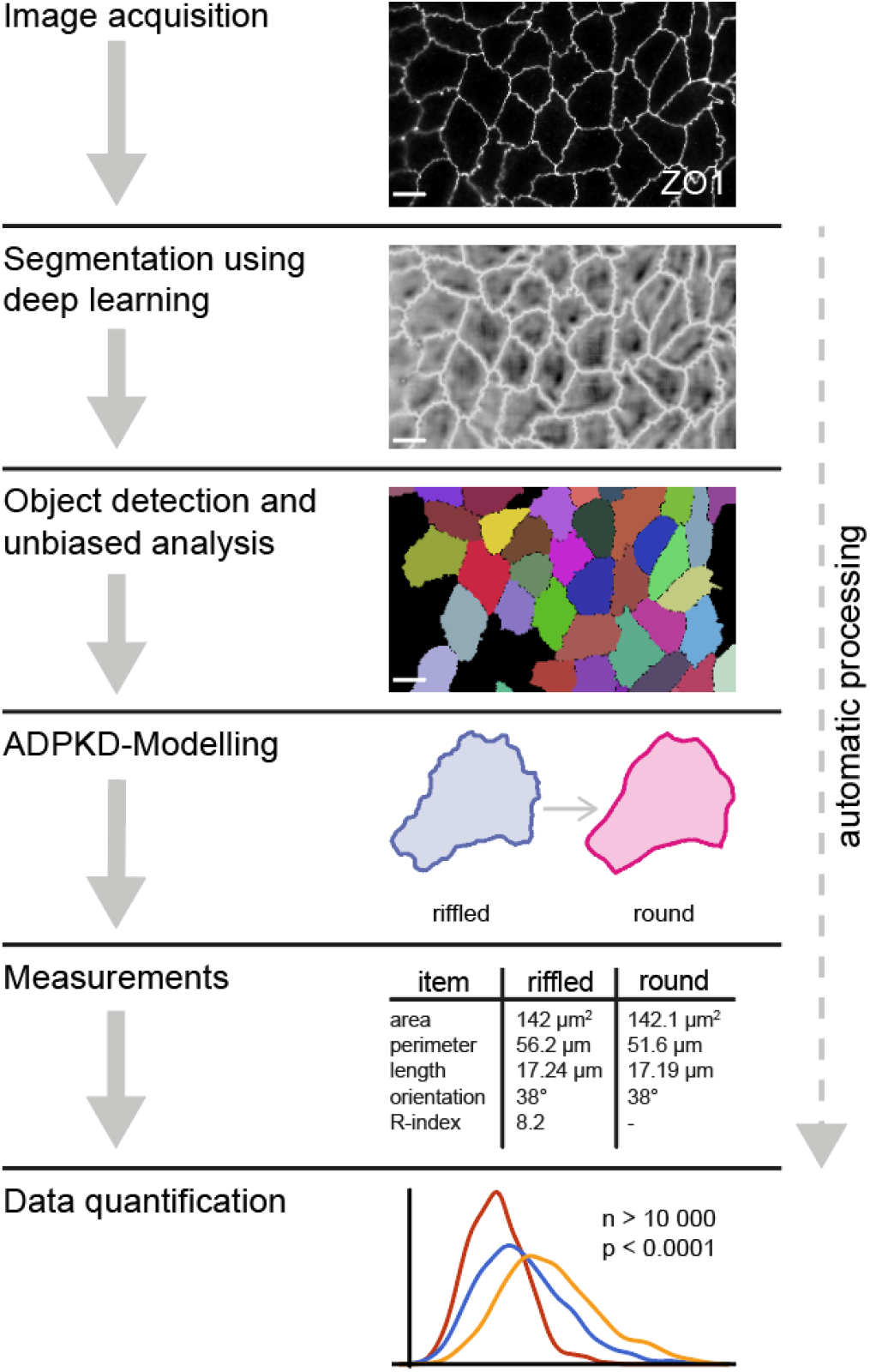
Schematic representation of the automated image analysis pipeline. Scale bars: 10 μm

### Cell detection and unbiased measurements

To ensure unbiased measurements, additional analysis steps were implemented in Mathematica 12 (Wolfram Research, Champaign, Illinois, USA). First, segmented images were pre-processed to remove background noise and eliminate incorrect segmentations. Tight junctions were then skeletonized to a single-pixel width to normalize variations in junction thickness observed across different genotypes and antibody combinations.

Watershed algorithms were used to detect individual cell objects from the skeletonized image (S3 Fig). This approach converts binary segmentations into meaningful cell-level data, capturing full cell geometries rather than only borders. We incorporated methods to ensure adequate segmentation and detection quality, retaining only cells with fully detected ZO1 staining and no contact with the image border. The error rate for false cell object detection was 0.4% (n=5.477). Various parametric values such as area, perimeter, and diameter were extracted for each object (Fig 3). All algorithms were standardized to ensure robust, reproducible measurements across datasets.

**Fig 3.**
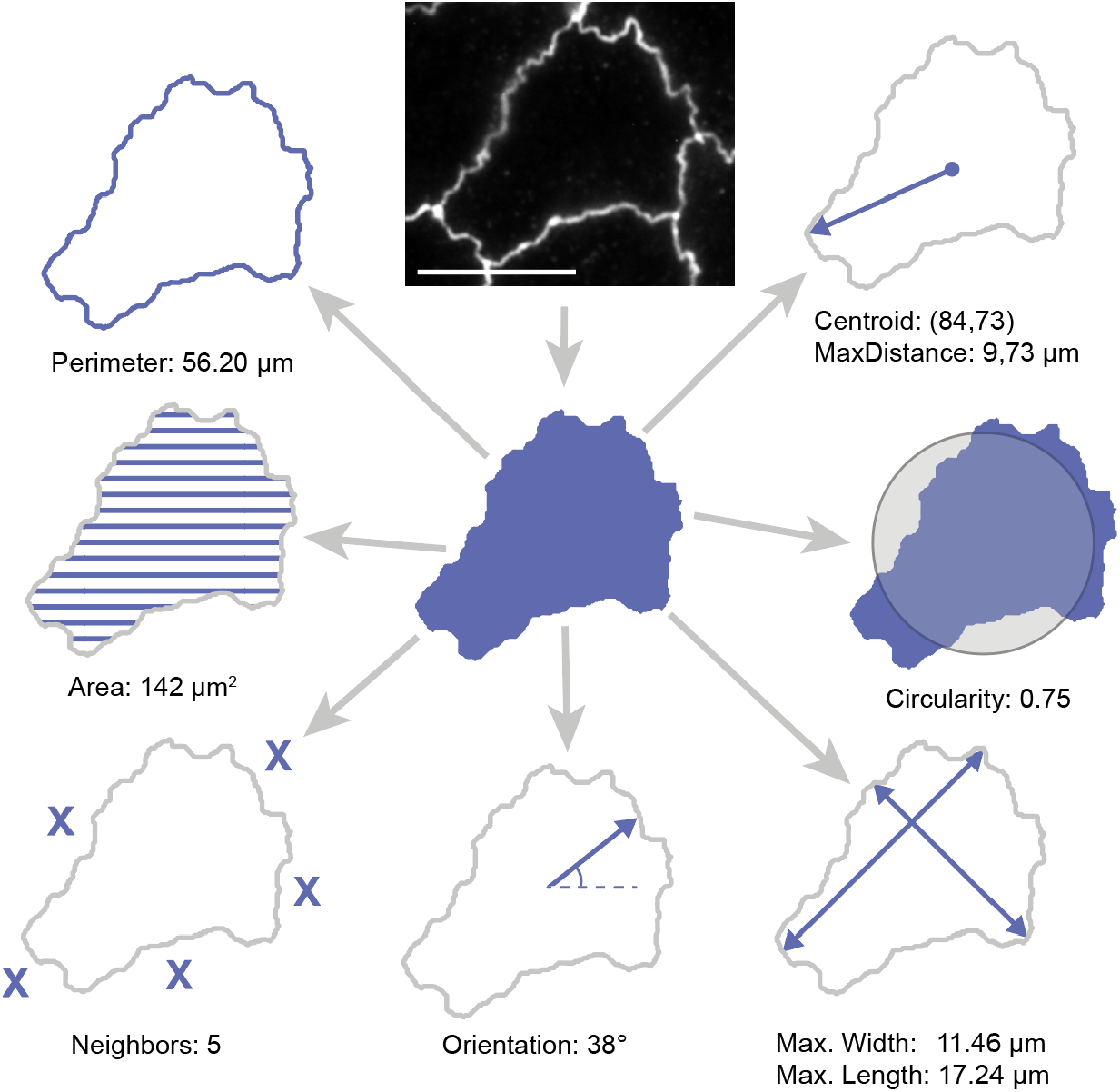
Visualization of potential measurement parameters extracted from cell objects. Scale bar: 10 µm

### ADPKD modeling and riffle pattern quantification

To better characterize the ADPKD phenotype and tight junction architecture, we developed a model to quantify membrane meandering and riffled patterns. While basic shape metrics such as circularity, a measure of object roundness (Fig 3), can be easily calculated, they did not reliably reflect the degree of tight junction irregularity in blinded manual assessments. Our approach addresses this limitation by comparing each segmented cell with a smoothed version of the original using a combination of blurring, erosion and edge filters (Fig 4A). By calculating and comparing the perimeters of the original and smoothed cell contours, we found that cells with pronounced riffles exhibit larger perimeter differences (Fig 4B). In contrast, cells with smoother, less meandering junctions had minimal differences between original and smoothened perimeters. The difference in perimeters was used to compute the riffle-index (R-index), a novel metric that quantifies tight junction irregularity (Fig 4C).

**Fig 4.**
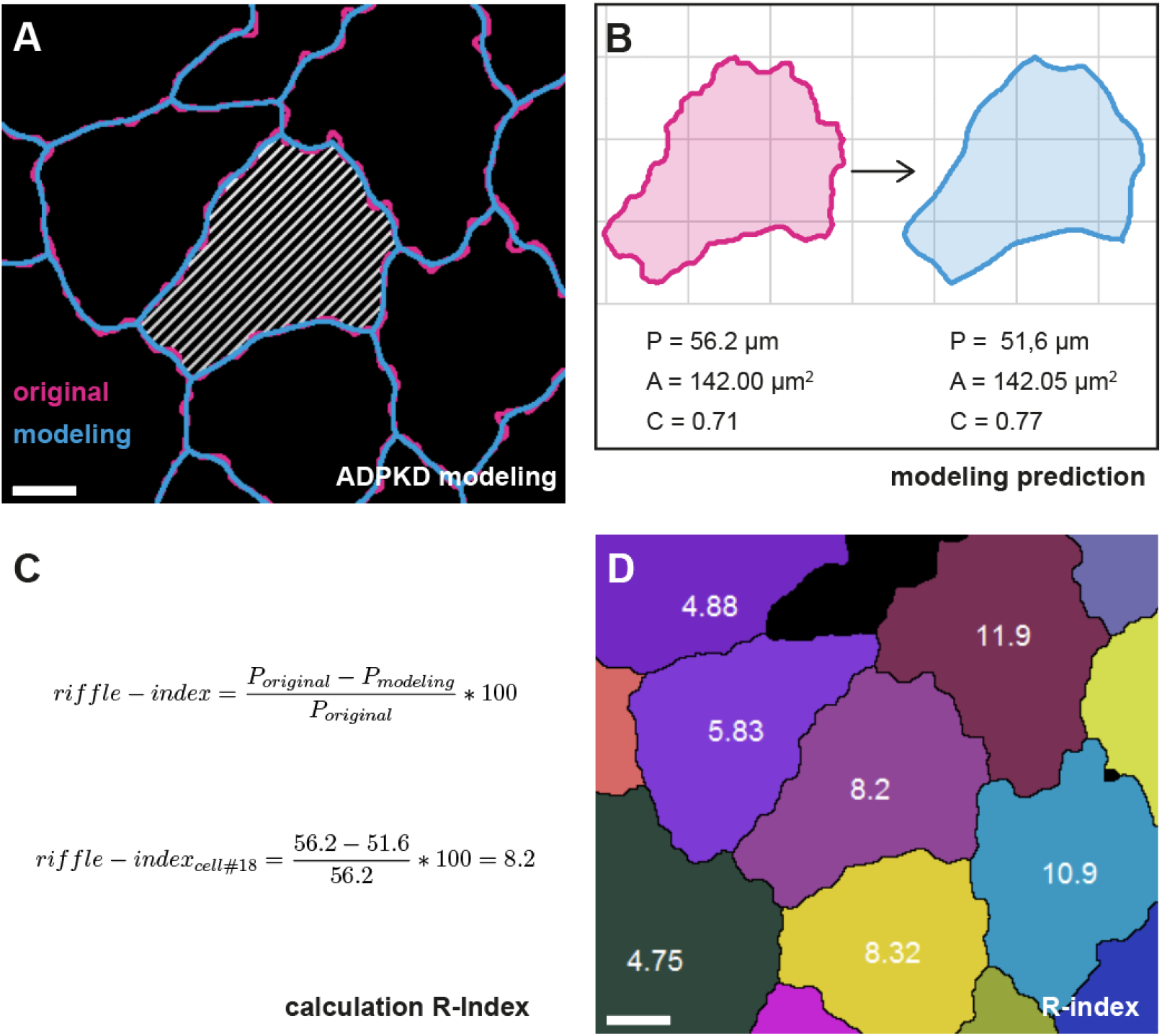
ADPKD modeling and computation of the R-index. (A) Overlay of the original segmentation (magenta) and the smoothed model (blue) of wild-type cells. Larger differences are observed when the original cell border is heterogeneous and displays pronounced ruffling. (B) Comparison of perimeter (P), area (A), and circularity (C) between the original and smoothed cell objects. (C) Illustration of R-index computation based on perimeter deviation. (D) Representative R-index values derived from individual cells. Scale bar: 4 μm.

Comparison of manual scoring with the computed R-index revealed strong concordance, with the automated analyses accurately reflecting visual assessments. The R-index reliably distinguished wild-type from *PKD2*^*-/-*^ cells and captured the rescue effect following *PKD2* re-expression (Fig 5). Moreover, the R-index enabled more precise, objective comparisons and statistical analysis across larger datasets and multiple genotypes than manual classification methods.

**Fig 5.**
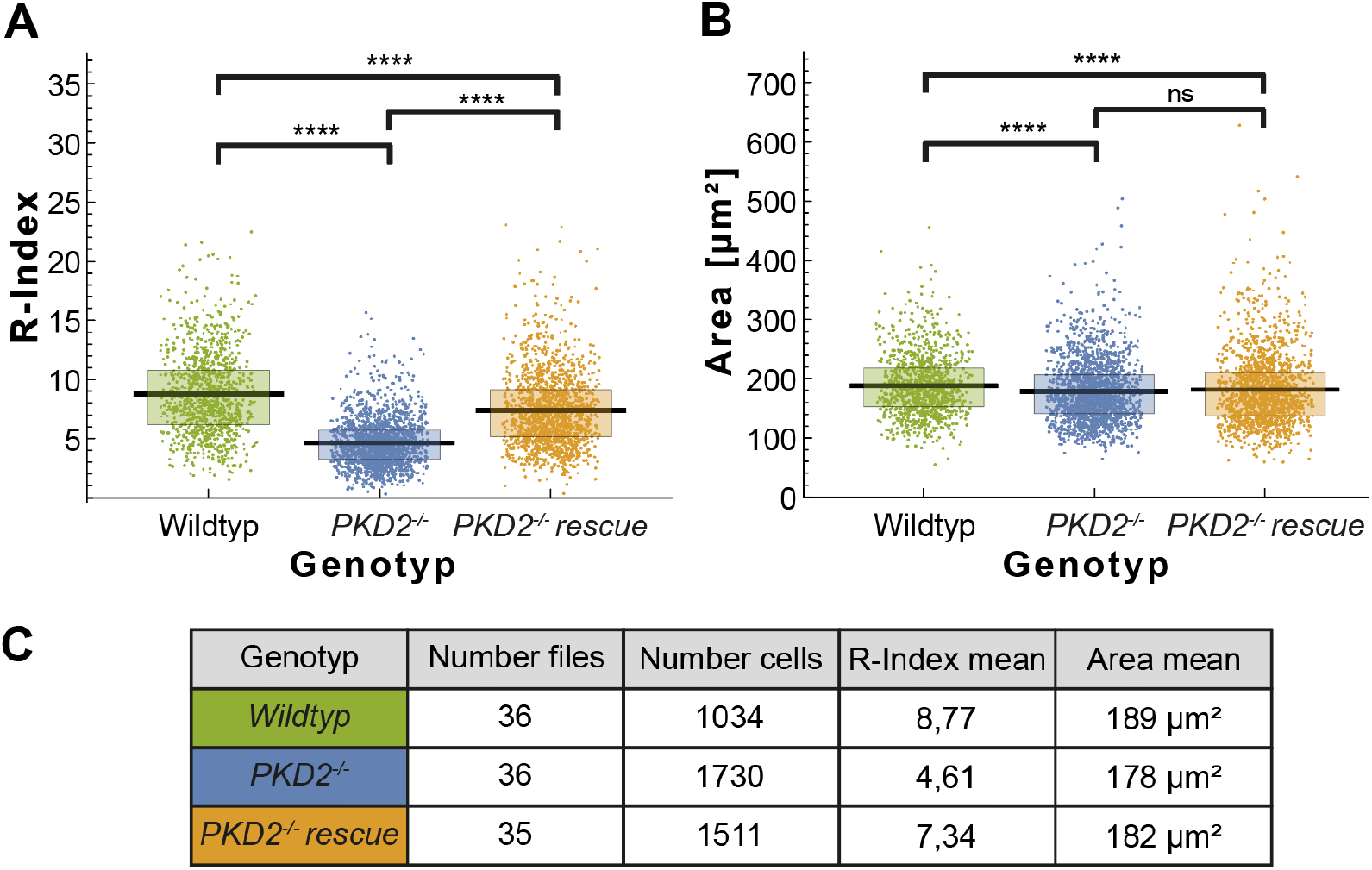
Data visualization in a high throughput setting. (A) Visualization of R-indices from a large image dataset. Each datapoint represents an individual cell. Mean value is shown by a black bar. The box includes the 25th to 75th percentile. (B) Same visualization for the cell area. (C) Summary statistics of the dataset shown. For both R-index and cell area, arithmetic means are reported. ns: not significant; *: p ≤ 0.05; ****: p ≤ 0.0001. Statistical testing was performed using the Mann–Whitney U test.

### Quantification of cytoskeletal traction between different phenotypes

To investigate the biomechanical mechanisms underlying riffled junction patterns and cell-cell interactions, we implemented models to estimate cytoskeletal traction at shared cell borders. We divided the segmented and skeletonized tight junctions into border segments formed between adjacent cell pairs. Traction was estimated by calculating the enclosed area of these segments (C in S4 Fig). Beyond monoculture experiments, this approach was extended to co-cultures containing GFP-labelled wild-type cells mixed with cells of another genotype. Phenotype-specific assignment of cell objects was achieved using an additional GFP fluorescence channel (A/B in S4 Fig). This framework may enable detection of alterations in cytoskeletal force transmission between cells of different genotypes, which are difficult to assess manually.

### Optimization for scalability and high throughput screening

The entire image processing pipeline was optimized for high-throughput screening. User interaction is minimal and limited to image selection, parameter specification, and quality control (Fig 2). We successfully processed datasets exceeding 3,000 images and more than 100,000 cell objects to screen a range of genotypes associated with the ADPKD spectrum. In addition to handling large datasets, individual image tiles were automatically stitched together to generate larger composite images (S5 Fig). This approach further reduces selection bias and enhances statistical power.

### Broader applications in cell biology

To evaluate the generalizability of the workflow, we applied it to immunofluorescence images of *Drosophila* egg chambers stained for b-catenin, commonly used to study cell shape regulation at the level of adherens junctions [20]. Without retraining, the segmentation algorithm achieved satisfactory performance. We then applied the same modeling algorithms to analyze membrane morphology, demonstrating the pipeline’s utility in studying cell shape regulation in other systems including model organisms *in vivo* (Fig 6).

**Fig 6.**
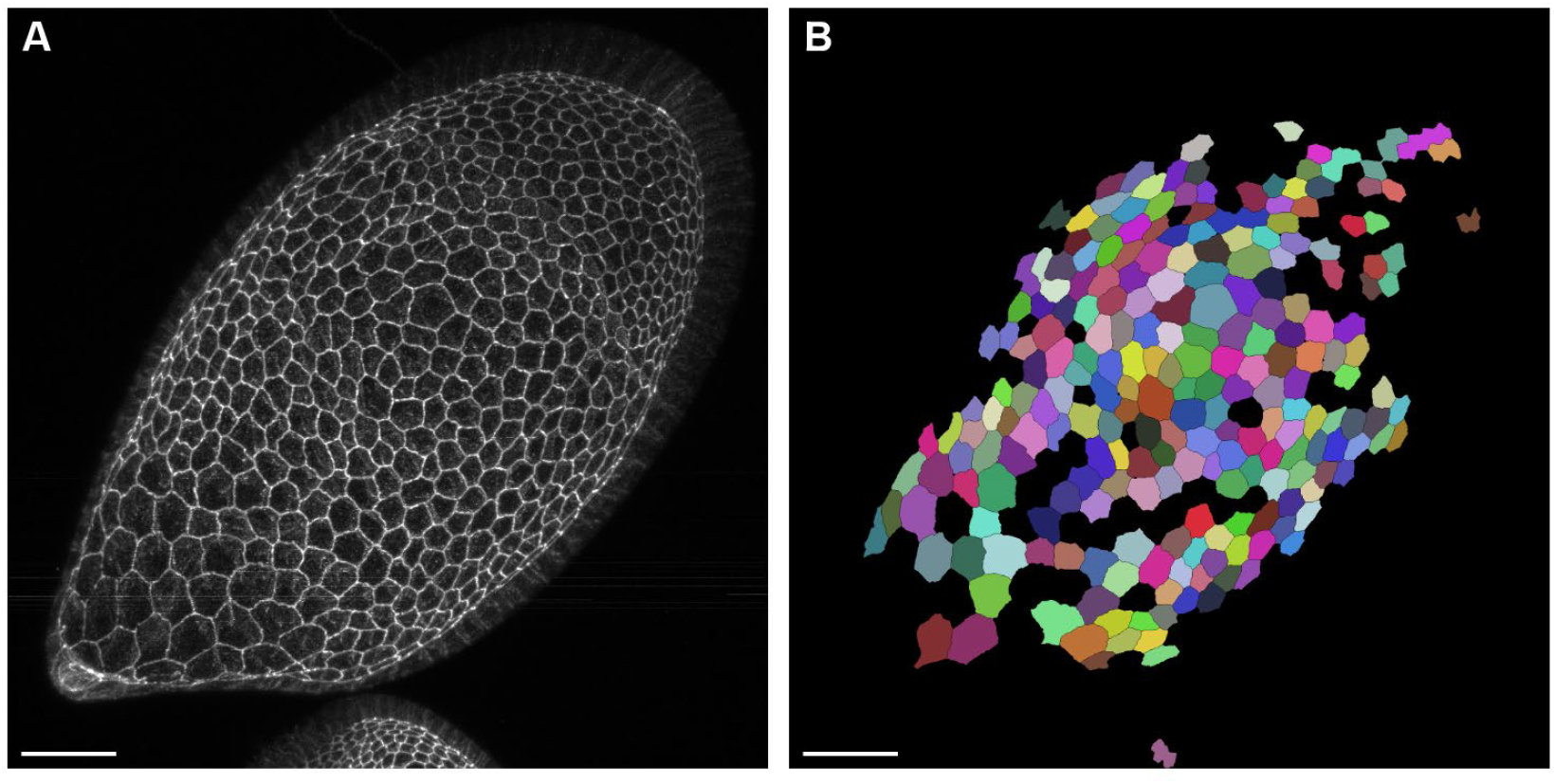
Application of the image analysis pipeline to dissect epithelial cell organization of Drosophila egg chambers. (A) Immunofluorescence image (Max. Proj.) of a *Drosophila* egg chamber stained for b-catenin, marking adherens junctions of the follicle epithelium. (B) Detected cells using the established pipeline, enable further quantification. Scale bars: 25 μm.

## Discussion

Cell shape is a fundamental determinant of tissue architecture and organ morphogenesis [21]. Epithelial cell geometry is shaped by cytoskeletal tension and junctional organization, which together govern organ development and function [1,3]. Disruption of the molecular mechanisms coordinating cell shape can contribute to pathological conditions, such as cystic kidney disease. In the context of ADPKD, cystogenesis is characterized by a shift from cuboidal to flattened cell morphology, a process that can be reversed through re-expression of *PKD* genes [22,23]. This highlights a connection between dysregulated cell shape and organ-level pathology. Understanding and quantifying epithelial morphology is therefore essential for elucidating disease mechanisms and developmental processes.

Despite this importance, the quantitative analysis of epithelial morphology remains challenging. Manual annotation of microscopy images is time-consuming, prone to user bias, and difficult to standardize, limiting reproducibility and scalability [24,25]. While traditional image analysis methods have provided valuable insights, they are often unsuitable for high-throughput applications or the resolution of subtle morphological details. Machine and deep learning-based approaches, offer scalable, standardized, and objective alternatives to overcome these limitations. U-Net architectures have proven especially effective for biomedical image segmentation [16,26] and are increasingly accessible via platforms such as ImageJ, enabling users with limited coding experience to train and apply segmentation models [15]. These systems offer flexibility and scalability, making them particularly valuable for analyzing large imaging datasets and subtle morphological phenotypes. Moreover, the reproducibility and standardization offered by deep learning models are crucial for building robust phenotypic profiles that can be compared across studies and platforms.

In this study, we developed a fully automated image analysis pipeline to quantify tight junction morphology in epithelial cells, with a focus on cell-based models for ADPKD. Our approach builds on the observation that ZO-1 stained MDCK cells exhibit distinctive junctional morphologies: wild-type cells display meandering and irregular borders, whereas *PKD1-* and *PKD2-*deficient cells show smooth, more linear borders. The partial restoration of the riffled phenotype upon *PKD2* re-expression supports the hypothesis that the gene product Polycystin-2 plays a critical role in maintaining epithelial morphology.

Using a U-Net-based segmentation of ZO1-stained monolayers, we extracted geometric cell features and introduced the R-index, a novel metric quantifying junctional meandering. The R-index captured phenotypic differences in a manner consistent with manual assessment, supporting its biological relevance. While minor systematic errors may arise from the multi-step modeling process, our automated approach clearly outperformed manual methods in terms of scalability, objectivity, and resolution of genotype-specific differences. Additional parameters, such as cell size and circularity, further enriched the morphological profiling allowed for a more comprehensive analysis.

Beyond general shape descriptors, we explored the mechanical basis of these morphological changes. We implemented a traction model analyzing curvature at shared cell borders. This approach decomposes junctional morphology into individual cell-cell interactions and can reveal alterations in force transmission, particularly in co-culture experiments where different genotypes may affect mechanical forces between neighboring cells. Such analyses are rarely feasible through manual inspection and offer a novel window into cellular mechanics in health and disease.

A major strength of our fully automated pipeline lies in its scalability and suitability for high-throughput screening. This is increasingly relevant in ADPKD research, where large-scale drug screens are being employed to identify pathogenic mechanisms and therapeutic candidates [27]. Compared to traditional image analysis, our automated pipeline reduces user-induced selection and confirmation bias and provides reproducible, quantitative measurements across large data sets. Nevertheless, systematic errors are an inherent risk of automated systems [28]. To mitigate these, we optimized U-Net training and modelling parameters to minimize general and systematic biases. Still, we recommend complementary manual inspection, particularly when applying the method to new cell types or imaging conditions. Ultimately, the modeling algorithms offer an approximation of biological reality, and these caveats must be recognized when interpreting results.

Limitations of the current approach should be acknowledged. Our analysis pipeline operates on 2D projections of epithelial monolayers and does not capture three-dimensional tissue architecture or interactions with extracellular matrix components known to affect cystogenesis in ADPKD [29]. Future iterations could incorporate 3D culture systems or organoids to improve the pathophysiological relevance of future analyses. Additionally, our traction estimates are derived from geometric proxies; direct biophysical measurements such as traction force microscopy would be valuable to validate our modeling approach [30].

We also demonstrated that a full end-to-end pipeline can be implemented using widely available tools such as ImageJ and Mathematica. While developing these systems requires coding knowledge, general-purpose platforms (e.g. CellProfiler [31]) offer accessible entry points for basic image analysis. Commercial solutions such as Aivia (Leica Microsystems, Wetzlar, Germany) also integrate deep learning, but may lack flexibility for specialized applications. For advanced experimental setups, custom solutions remain necessary.

In summary, we present a deep learning-based workflow for the automated quantification of epithelial cell shape and tight junction morphology. The integration of junctional analysis with traction modeling provides novel tools for investigating cytoskeletal regulation and epithelial morphogenesis. These methods, combined with genetically modified cell models and standardized imaging protocols, enable robust, scalable, and reproducible phenotyping applicable to a wide range of biological questions. Our findings highlight the power of automated image analysis to move beyond manual inspection, providing deeper, data-driven insight into the complex interplay between cell shape, mechanics, and function in epithelial biology, with implications for disease pathogenesis and morphogenetic regulation.

## Materials and Methods

### Cell lines

Madin Darby Canine Kidney cells (MDCK) (American Type Culture Collection, Manassas, USA) as well as genetically modified *Pkd1*^*-/-*^, *Pkd2*^*-/-*^, *Pkd2*^*-/-*^ rescue, Cgn^*-/-*^, and WT.GFP lines, were cultivated as adherent monolayers in Dulbecco’s modified Eagle’s medium (Thermo Fisher Scientific Inc., Waltham, USA) supplemented with 10% heat-inactivated fetal bovine serum (Sigma-Aldrich, St. Louis, USA). Cell lines were maintained in a humidified incubator at 37°C with 10% CO_2_ and passaged every 2-3 days using 0.25% trypsin-EDTA (Thermo Fisher Scientific Inc., Waltham, USA).

### Immunofluorescence staining

Immunofluorescence staining of MDCK cells was performed as previously described [32]. Briefly, cells were seeded on cover glasses (Carl Roth) in six-well plates (Greiner Bio-One, Germany) and grown for ten days. The α-ZO1 primary antibodies (Santa Cruz Biotechnology, Dallas, USA and Thermo Fisher Scientific Inc., Waltham, USA) were diluted 1:50 or 1:500 in PBS, respectively. Imaging was performed on an Axio Observer.Z1 microscope (Zeiss, Jena, Germany) equipped with a Zeiss i-Plan-Apochromat 63x/1.40 Oil DIC M27 objective using ZEN 2 (blue edition) Version 2.0.0.0. Pixel size: 0.102µm/px. Tile functions were used to generate stitched images.

### *Drosophila* stocks and genetics

*Drosophila melanogaster* stock *w[118]* was used. Stock was maintained on standard fly food (10 L water, 74.5 g agar, 243 g dry yeast, 580 g corn meal, 552 mL molasses, 20.7 g Nipagin, 35 mL propionic acid) at 22 °C. Flies were fed yeast paste for 48 to 72 h prior to dissection.

### Immunohistochemistry and imaging of *Drosophila* egg chambers

Ovaries were dissected and fixed in 4% paraformaldehyde/PBS for 15 min at 22°C. Washes were performed in PBS + 0.1% Triton X-100 (PBT). Samples were incubated with primary antibody mouse anti-β-catenin (1:100, DSHB, N27A1) in PBT overnight at 4°C. Ovaries were washed with PBT and then incubated with dyes and secondary antibodies for 2 h at 22 °C. DAPI (1:500, Sigma), Phalloidin (1:500 Alexa Fluor 488) was used to visualize DNA and filamentous Actin, respectively. Donkey anti-mouse Alexa Fluor647 (Abcam, AB150111, 1:500) was used as secondary antibody. Samples were mounted using Molecular Probes Antifade Reagents and imaged using the Leica TCS SP8 confocal microscope.

### Training the deep learning network

The U-Net model was trained using the ImageJ U-Net plugin [15]. Image annotation was conducted in 3D Slicer [33] following the general instructions provided by Falk et. al. [15]. These labelling masks were converted to ImageJ (U. S. National Institutes of Health, Bethesda, Maryland) regions-of-interest to match the plugin requirements. Training used N = 9 partly labelled images with the following settings: „Training from scratch” with „2D Cell Net (v0)”-model; element size: 0.102 µm x 0.102 µm; Input patch size: 1020×1020 px; learning rate: 10E-4, iterations: 10 000. Computation was performed on a remote server running Ubuntu 16.04.6 LTS (Canonical, London, UK) and the precompiled U-Net library with a Nvidia (Santa Clara, California) Geforce GTX 1080 Ti with 12GB GDDR5 RAM.

### Image analysis

A custom Fiji ImageJ (version 1.53b) script was developed for automatic pre-processing and segmentation enabling high throughput analysis. Subsequent analysis steps were implemented in Mathematica 12.0. The source code, sample data and documentation are available on GitHub at https://github.com/JahnJ/U-Net-based-image-quantification-of-epithelial-cell-shapes/. We have also used Zenodo to assign a DOI to the repository: 10.5281/zenodo.15857792.

## Acknowledgements

The authors acknowledge Simone Diederichsen for expert technical assistance. We would like to thank the Life Imaging Center (LIC) of the University Freiburg for their assistance with server hosting.

