## Supplemental Figure 3 for "Deep learning-based image quantification of epithelial cell shapes and its application to polycystic kidney disease"

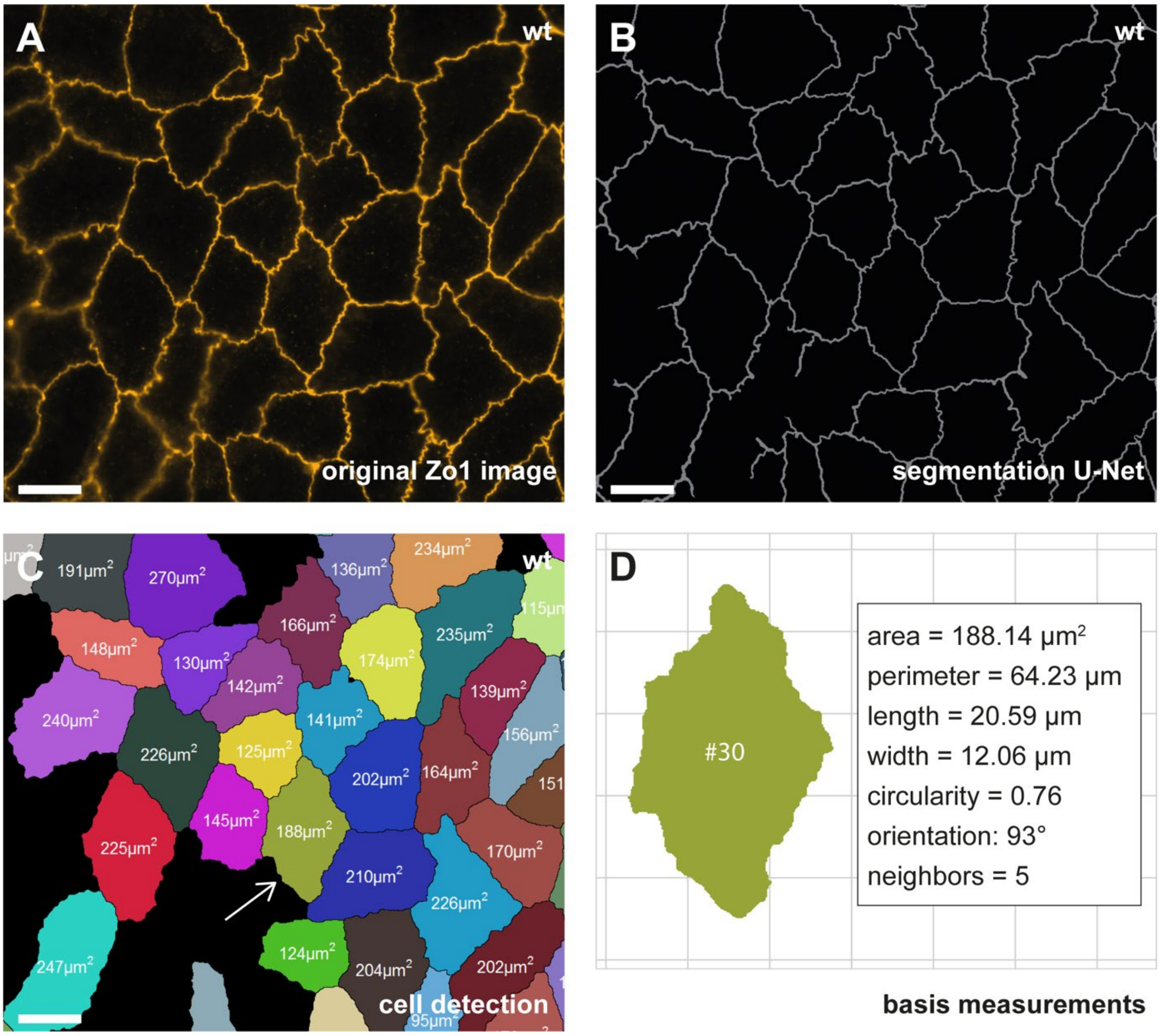


**S3 Fig. Detection of individual cell objects from tight junction staining.**

(A) MDCK cells were fixed, stained with ZO-1 antibodies, and imaged using immunofluorescence microscopy to visualize tight junctions. (B) The trained U-Net deep learning model was applied to segment cell borders. Regions with insufficient signal or out-of-focus areas were excluded to maintain segmentation accuracy. (C) Individual cell objects were extracted from the segmentation using an adapted watershed algorithm. Area values are shown for each detected object. The full sample contains 45 detected cells - a cropped region is shown for improved visibility. (D) Multiple morphological parameters, such as area and perimeter, can be calculated for each cell object. Scale bar: 10 µm (A–C). MDCK = Madin-Darby Canine Kidney;  ZO1 = Zonula occludens protein 1.
