## Supplemental Figure 4 for "Deep learning-based image quantification of epithelial cell shapes and its application to polycystic kidney disease"

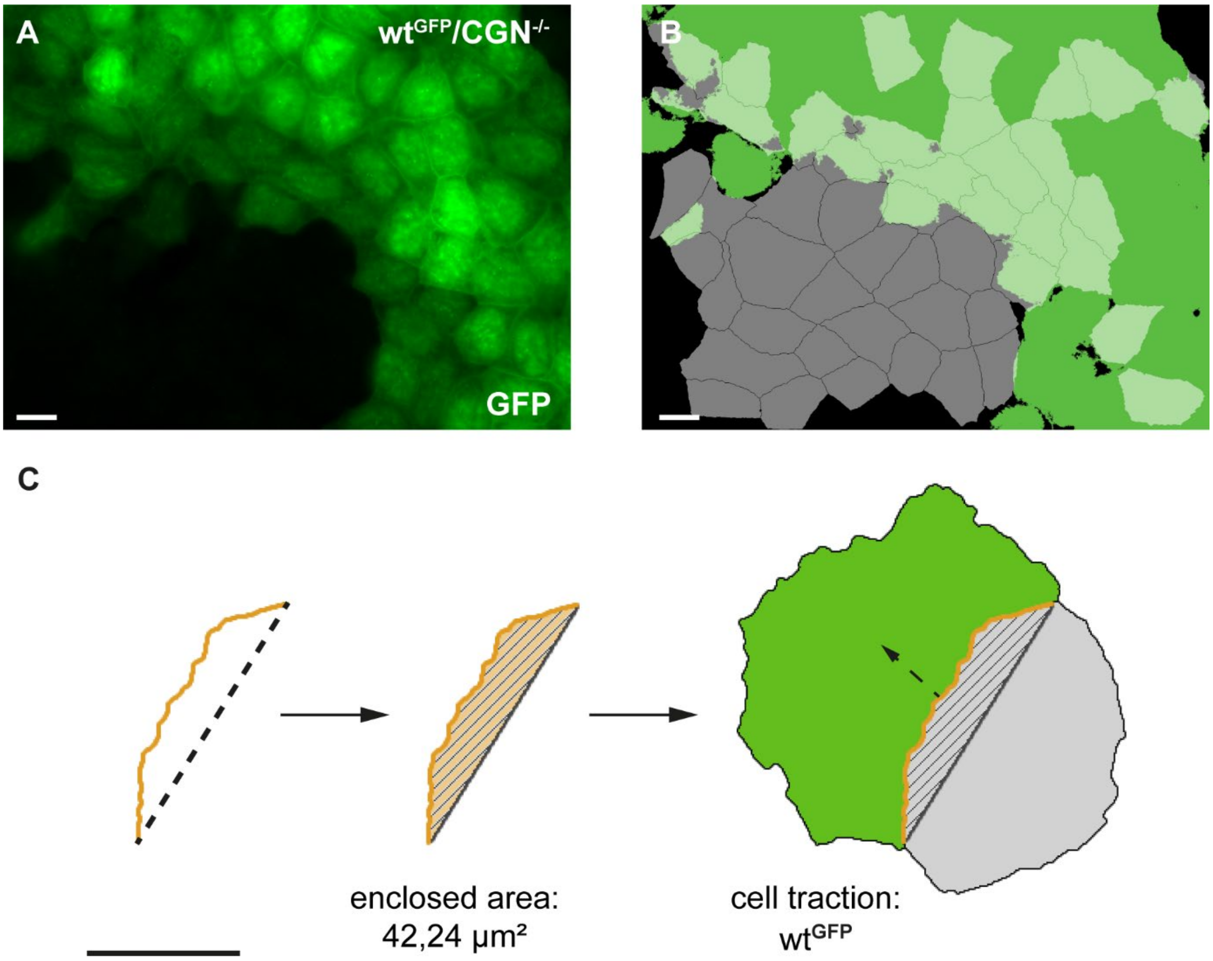


**S4 Fig. Genotype-specific cell interaction analysis in mixed cultures.**

(A) In a mixed culture of GFP-labeled WT and unlabeled $CGN^{-/-}$ MDCK cells, GFP intensity was used to identify WT cells. (B) Overlay of the GFP signal with segmented cell objects allowed classification of each cell as “GFP-positive” (WT) or “GFP-negative” ($CGN^{-/-}$) based on spatial overlap. Loss of the tight-junction-associated protein Cingulin phenocopies loss of Polycystin-2. (C) Example of two neighboring cells of different genotypes. The WT cell is shown in green, the $CGN^{-/-}$ cell in gray. The enclosed area between their junctions was calculated from segment endpoints and used to quantify traction. In this case, traction is directed toward the wild-type cell. Scale bar: 10 µm. Cgn = Cingulin; GFP = Green fluorescent protein; WT = Wildtype.
