## Supplemental Figure 2 for "Deep learning-based image quantification of epithelial cell shapes and its application to polycystic kidney disease"

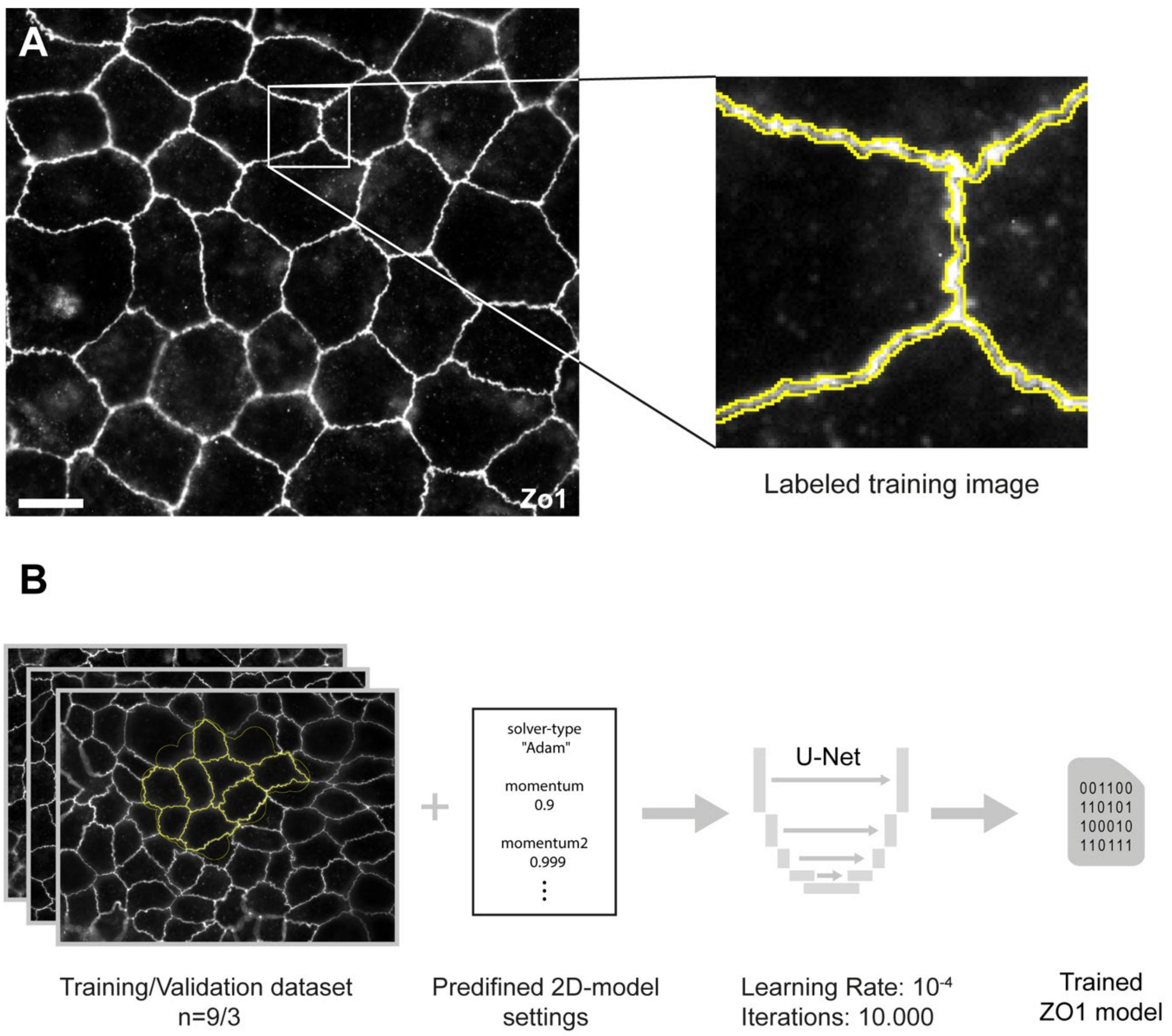


**S2 Fig. Training procedure for the ZO1 deep learning model.**

(A) Representative immunofluorescence image with manually annotated tight junction structures used for training. (B) Annotated images were split into a training and validation dataset and used to train a U-Net convolutional neural network. The displayed parameters (learning rate and number of iterations) can be adjusted as needed. Once trained, the ZO1 segmentation model can be applied to new image data for automated analysis. Panel (B) is adapted from [15]. Scale bar: 10 µm. ZO1 = Zonula occludens protein 1.
