## Supplemental Figure 5 for "Deep learning-based image quantification of epithelial cell shapes and its application to polycystic kidney disease"

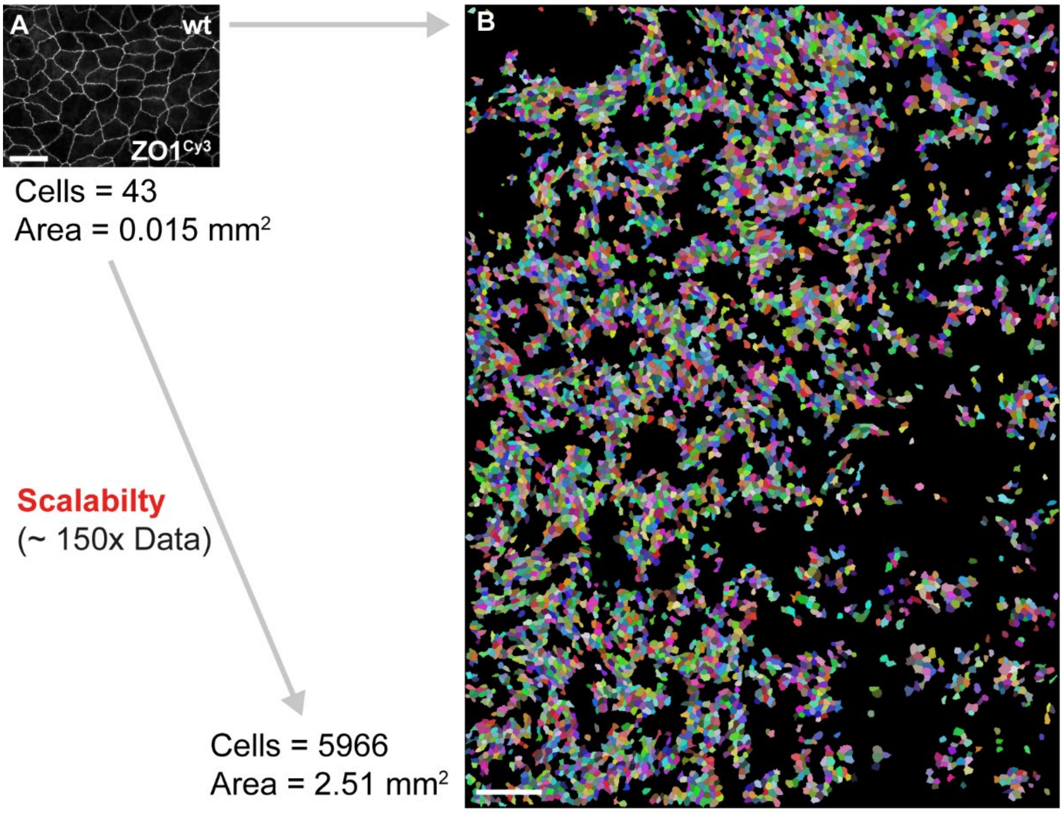


**S5 Fig. Scalability for dataset expansion through image stitching.**

Immunofluorescence images of MDCK WT cells stained for ZO-1 are shown. Panel (A) displays a single image acquired with a 63× objective using a Cy3-labeled secondary antibody. Panel (B) shows a larger, automatically stitched image generated from a 9×9 tile scan using a 40× objective and an Alexa Fluor 488 labeled secondary antibody. The stitching process, performed directly at the microscope, enabled a substantial increase in image area and data volume. Total processing time for the stitched image was approximately 3 hours. Larger black regions in (B) indicate areas with insufficient staining or out-of-focus regions that were excluded from analysis. Scale bars: 25 µm (A), 150 µm (B). MDCK = Madin-Darby Canine Kidney; ZO1 = Zonula occludens protein 1; WT = Wildtype.
