## Supplemental Figure 1 for "Deep learning-based image quantification of epithelial cell shapes and its application to polycystic kidney disease"

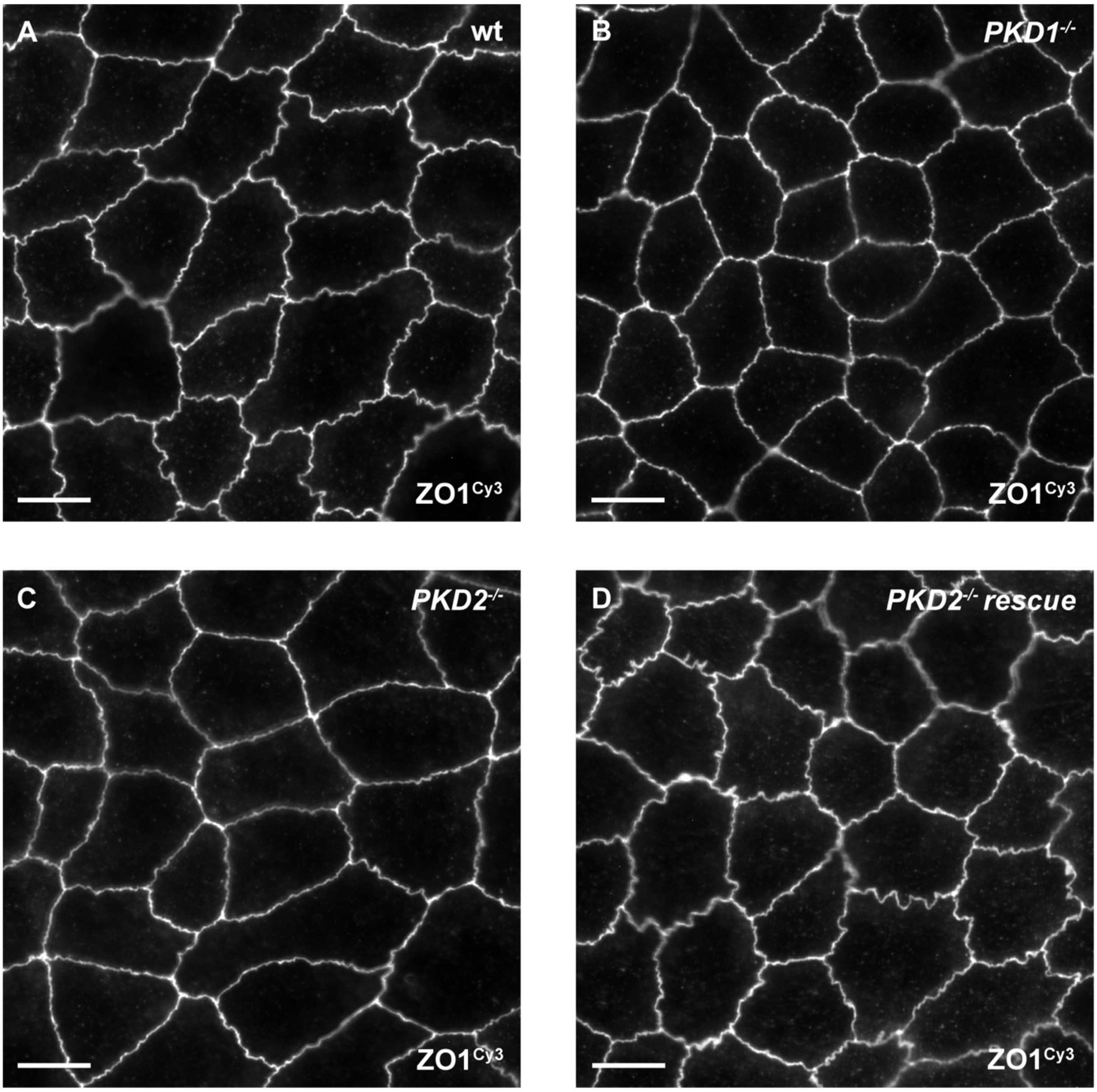


**S1 Fig. Meandering in ADPKD-related genotypes.**

Representative immunofluorescence images of MDCK WT (A), *PKD1*$^{-/-}$ (B), *PKD2*$^{-/-}$ (C) and *PKD2*$^{-/-}$ *rescue* (D) cells, which were stained with an α-Zonula occludens 1 primary antibody and a Cy3-conjugated secondary antibody. (A) WT cells show prominent membrane riffles. (B/C) *PKD1*$^{-/-}$ and *PKD2*$^{-/-}$ cells exhibit smooth cell borders with minimal indentations. (D) Re-expression of *PKD2* restores the meandering morphology of tight junctions. Scale bar: 10 µm. MDCK = Madin-Darby Canine Kidney; *PKD1* = Polycystic kidney disease gene 1; *PKD2* = Polycystic kidney disease gene 2; WT = Wildtype; ZO1 = Zonula occludens protein 1.
